## Supplemental methods and figures for "Quantifying yeast microtubules and spindles using the Toolkit for Automated Microtubule Tracking (TAMiT)"

### Supplemental Material

#### 1 Feature Inheritance

TAMiT implements features as instances of classes. The structure of inheritance is shown in Fig. **??**. The most general class is a *Feature*. There are two types of features. *Basic Elements* are physical and geometric objects that are associated with fluorescent intensity. These include lines, spots, and curves. *Organizers* structurally organize the basic elements to form composite features. An organizer can contain basic elements or other organizers. For example, an *Aster* contains one spot, and multiple lines. A *Spindle* and a *Spindle Bud* contains a line (for the spindle), and two asters (for SPB and astral microtubules).

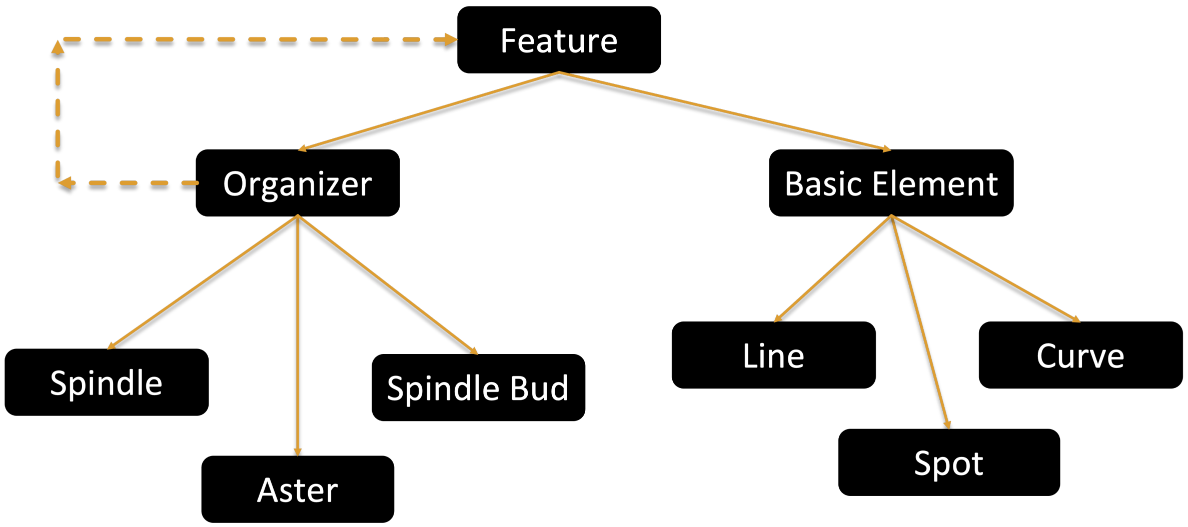


Figure S1: Inheritance in TAMiT. A feature is the most general object. There are two kinds of features, an Organizer and a Basic Element. Basic Elements are physical and geometric objects that have an intensity profile. Organizers are composed of other features and organize them in various ways. For example, the aster organizer contains a spot and multiple lines.

#### 2 SPB detection

The detection process is independent for each frame in the image stack. For a 3D image (XYZ), we take a maximum intensity projection along Z. This gives us a 2D image, which we use for the estimation process. Here, we show the detection process for a spindle pole body (SPB) in a *S. pombe* monopolar spindle (Fig. S2A). We begin with pre-processing where we apply a gaussian filter on the image (Fig. S2B). This gaussian filter (standard deviation equal to 2 pixels) is used to enhance the tubulin signal. The next step is to identify the region of the SPB. We perform otsu thresholding, followed by an extended-maxima H transform to find an extended region belonging to the SPB (Fig. S2C). Finally, the maximum intensity pixel within this region is used as the estimate for the SPB location (Fig. S2D). For the Z position of the SPB, we find the Z pixel, with fixed XY location, with the maximum intensity.

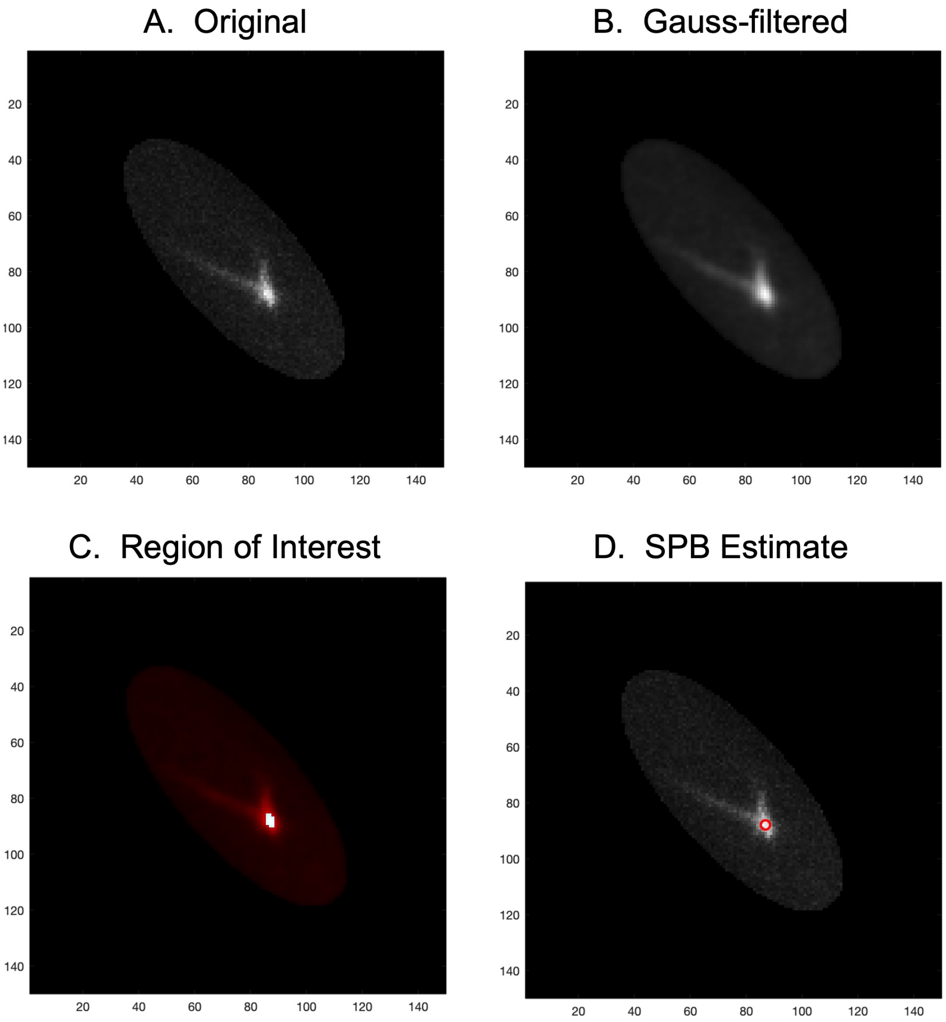


Figure S2: Spindle pole body (SPB) detection. All images are maximum intensity projections in Z of the full 3D image. (**A**) Original image showing a monopolar spindle. The SPB is the brightest region. (**B**) Gaussian-filtering with a filter of size equal to the point spread function enhances the fluorescent signal of tubulin. (**C**) Otsu thresholding followed by an extended maxima transform yields a region of interest. (**D**) SPB estimate is the maximum intensity pixel in the region of interest.

### 3 Straight microtubule detection

Line detection for straight microtubules starts with an estimation for a start point. This may be done using either SPB detection for a *S. pombe* monopolar spindle (Sec. 2) or spindle detection for a *S. pombe* mitotic spindle (Sec. 4). The line algorithm executes the following steps:

1. Given a start point (Fig. S3A), perform an azimuthal sweep and calculate the summed intensity along different directions.

2. Find peaks in the intensity plot to identify angles with enhanced intensities (Fig. S3B).

3. For each identified peak/angle, measure the intensity at different radii (Fig. S3C).

4. We smooth the signal and find the radius when it falls below a threshold $T_1$, defined as one standard deviation above the image background ($T_1 = \mu + 1\sigma$) (Fig. S3C). This threshold value

can be increased to multiple standard deviations if the signal to noise ratio of the image is greater. This can be used to find the length of the line.

5. Given the start point, an angle, and a length, we find the end-point of each line.

The final estimate is shown in Fig. S3D.

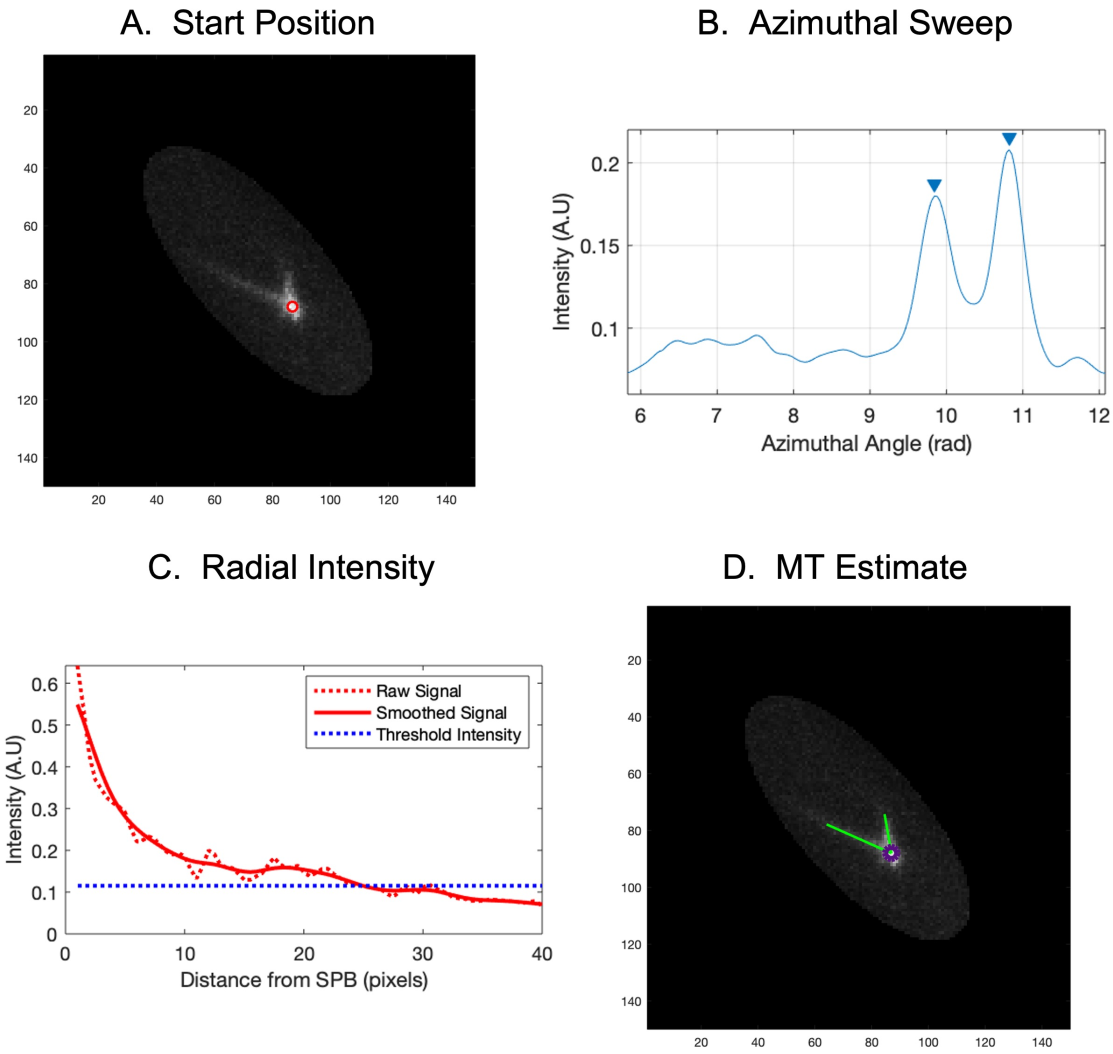


Figure S3: Detection of straight microtubules. All images are maximum intensity projections in Z of the full 3D image. (**A**) The start position where the microtubule search algorithm is initiated. This is normally the SPB location. (**B**) Azimuthal sweep gives the summed intensity at different angles. A peak finder gives two clear peaks corresponding to the two microtubules visible in the original image. (**C**) Radial intensity along the direction of one of the microtubules. A threshold determines the end-point of the microtubule along this direction. (**D**) The final estimate showing the detected microtubules in green and the SPB in purple.

#### 4 Spindle detection

We start with a 2D maximum intensity projection in Z as mentioned earlier, and apply a gaussian filter (standard deviation of 2 pixels) (Fig. S4A). Next, we use iterative thresholding to find an elongated region in the image. The thresholding process works as follows:

1. Initialize a threshold value equal to the median intensity of the image

2. Generate a boolean mask with pixels above the threshold given value 1 and below the threshold given value 0.
3. Find a convex hull (ensures that there is a single connected region in the image
4. Find the major and minor axis length of this region, and subsequently the aspect ratio.
5. If the aspect ratio is less than 4 (tunable parameter, should be set around 5 to yield a region that represents the spindle), then increase threshold by 1% and repeat steps 2-5. If condition is satisfied, or a maximum threshold value is reached, then stop iterative thresholding and accept the boolean mask (Fig. S4B).

Given the mask (which may contain multiple regions), pick the region with the greatest integrated intensity. Next, we find the corresponding major axis which is expected to align with the spindle axis (Fig. S4C). Measuring the intensity along this axis, we find two end points where the normalized intensity drops to 0.6 (Fig. S4D). This threshold can be increased for cases where the signal to noise ratio is higher and there is no considerable intensity variation within the spindle. The ratio can be lowered in cases where there is significant intensity variation within the spindle, either in the middle or at the ends. These end-points correspond to the SPB locations and the line connecting them represents an estimate for the spindle (Fig. S4E).

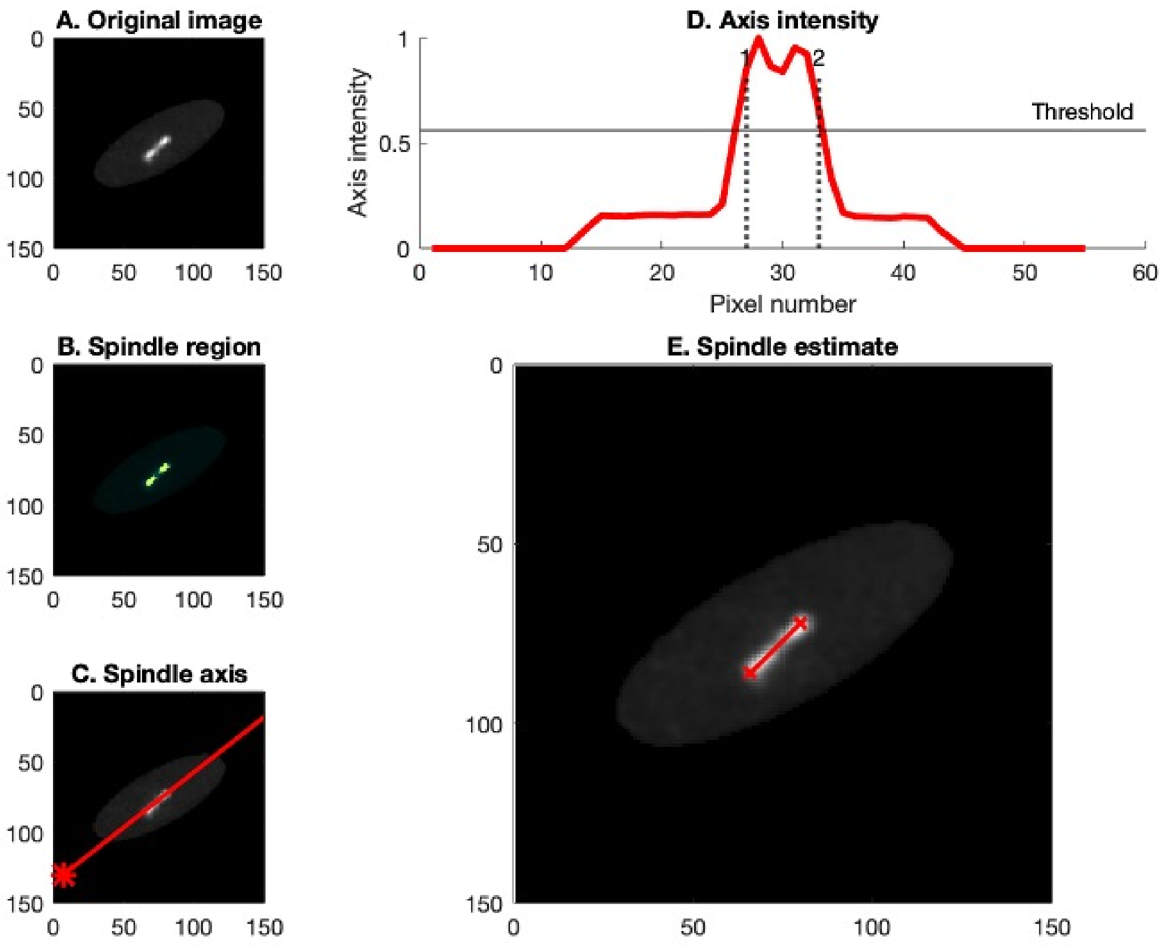


Figure S4: Spindle detection. All images are maximum intensity projections in Z of the full 3D image. (**A**) Original image showing the bright spindle. (**B**) Iterative thresholding yields regions of interest belonging to the spindle. (**C**) Axis passing through the best region of interest. (**D**) Intensity along the axis. A threshold determines the two end-points belonging to the the spindle end-points. (**E**) The spindle estimate.

### 5 Curved microtubule detection

Here, we show curve detection performed on a time frame from *S. cerevisiae* (Fig. S5A). To find curved microtubules, we need estimated start points (Fig. S5B). We begin by using steerable filtering to find pixels showing line-like intensity (Fig. S5C). This gives us paths which can be identified using iterative algorithms. For a given start point, we begin by performing an azimuthal sweep to identify the number of curves and the initial direction of curves emanating from this point. Then, for each curve, we find its coordinates by following these steps:

1. Begin with a start point, an initial angle, and a path image that comes from steerable filtering (Fig. S5C).

2. Find the next point along the path by performing an azimuthal scan, finding the highest intensity peak, and taking a small radial step in that direction.

3. Continue until the highest intensity peak is below a threshold value $T_2$. This value is set equal to the mean intensity of the path image (background value) and does not need to be altered. This background value signifies the general line-like signal in the path image, and pixel values must be above this background for them to signify the presence of a line.

4. Reduce the radial step size and attempt to find another point. If intensity is still below threshold $T_2$, reject and stop finding more points along this path/curve. If intensity is now above $T_2$, repeat steps 2 and 3 until no more points are found.

5. This concludes the path finding algorithm. The final coordinates are shown in Fig. S5D,E.

6. Now, using the original image (instead of the path image), find the intensity at the microtubule end point. If the intensity is below threshold $T_1$ (sum of image median and one standard deviation), reduce the length of found curves until the intensity at the end-point satisfies the threshold criterion.

The final estimate with curved microtubules is shown in Fig. S5F.

### 6 *S. pombe* Strains

### 7 Movies

- Movie 1: Microtubule detection in *S. pombe* mitotic cell. Left (original MIP), center (fitted MIP), and right (2D features). Colorbar represents the position in Z ($\mu m$).

- Movie 2: Microtubule detection in *S. pombe* monopolar cell. Left (original MIP), center (fitted MIP), and right (2D features). Colorbar represents the position in Z ($\mu m$).

- Movie 3: Microtubule detection in *S. cerevisiae* mitotic cell. Left (original MIP), center (fitted MIP), and right (2D features). Colorbar represents the position in Z ($\mu m$).

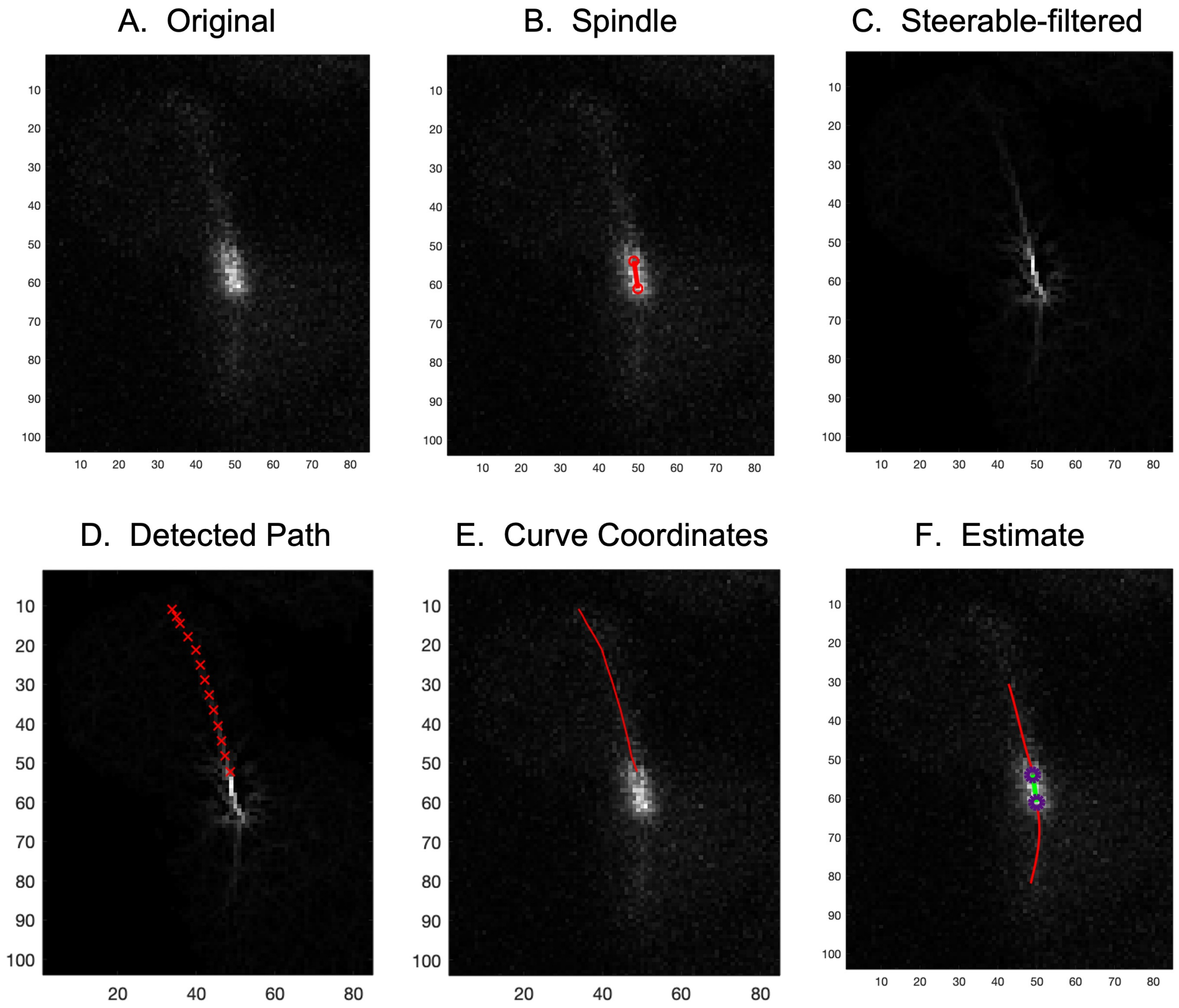


Figure S5: Detection of curved microtubules. All images are maximum intensity projections in Z of the full 3D image. (**A**) Original image showing a budding yeast metaphase cell. (**B**) The detected spindle found using the algorithm presented in Sec. 4. (**C**) Steerable filtering accentuates pixels where a line passes through. (**D**) Detected path of a curve (red) using a combination of a restricted azimuthal sweep, and a fixed radial step. (**E**) Curve coordinates generated from the path shown in D. (**F**) Final estimate of the curved microtubules (red) along with the spindle (green) and the SPBs (purple). Microtubule length is reduced based on intensity considerations mentioned in Sec. 5.

| Strain # | Genotype | Source |
|---|---|---|
| MB850 | cut7-24, Z:adh15:mCherry-atb3:natMX6, ura4-D18, h+ | this study |
| MB857 | cut7-24, Z:adh15:mCherry-atb3:natMX6, klp6Δ::ura4+, ura4-D18, h- | this study |
| MB863 | cut7-24, Z:adh15:mCherry-atb3:natMX6, alp14Δ::Kanr, ura4-D18, h+ | this study |
| MB890 | cut7-24, Z:adh15:mCherry-atb3:natMX6, cls1-36:ura4+, ura4-D18, h+ | this study |
| MB919 | cut11-7, Z:adh15:mCherry-atb3:natMX6, mis12-GFP::leu1+, leu1-32, ura4-D18, h+ | this study |

Figure S6: S. pombe strain information
